# Global analysis of the RpaB regulon based on the positional distribution of HLR1 sequences and comparative differential RNA-Seq data

**DOI:** 10.1101/443713

**Authors:** Matthias Riediger, Taro Kadowaki, Ryuta Nagayama, Jens Georg, Yukako Hihara, Wolfgang R. Hess

**Affiliations:** Genetics & Experimental Bioinformatics, Institute of Biology III, Faculty of Biology, University of Freiburg, Schänzlestr. 1, 79104 Freiburg, Germany; Graduate School of Science and Engineering, Saitama University, Saitama 338-8570, Japan; Freiburg Institute for Advanced Studies, University of Freiburg, Albertstr. 19, D-79104 Freiburg, Germany

**Keywords:** cyanobacteria, OmpR-type transcription factors, photosynthetic gene expression, regulatory networks, sRNAs

## Abstract

The transcription factor RpaB regulates the expression of genes encoding photosynthesis-associated proteins during light acclimation. The binding site of RpaB is the HLR1 motif, a pair of imperfect octameric direct repeats, separated by two random nucleotides. Here, we used high-resolution mapping data of transcriptional start sites (TSSs) in the model *Synechocystis* sp. PCC 6803 in conjunction with the positional distribution of HLR1 sites for the global prediction of the RpaB regulon. The results demonstrate that RpaB regulates the expression of more than 150 promoters, driving the transcription of protein-coding and non-coding genes and antisense transcripts under low light and upon the shift to high light when DNA binding activity is lost. Transcriptional activation by RpaB is achieved when the HLR1 motif is located 66 to 45 nt upstream, repression occurs when it is close to or overlapping the TSS. Selected examples were validated by multiple experimental approaches, including chromatin affinity purification, reporter gene, northern hybridization and electrophoretic mobility shift assays. We found that RpaB controls *ssr2016/pgr5*, which is involved in cyclic electron flow and state transitions; six out of nine ferredoxins; three of four FtsH proteases; *gcvP/slr0293*, encoding a crucial photorespiratory protein; and *nirA* and *isiA* for which we suggest cross-regulation with the transcription factors NtcA or FurA, respectively. In addition to photosynthetic gene functions, RpaB contributes to the control of genes affiliated with nitrogen assimilation, cofactor biosyntheses, the CRISPR system and the circadian clock, making it one of the most versatile regulators in cyanobacteria.

**Significance Statement:** RpaB is a transcription factor in cyanobacteria and in the chloroplasts of several lineages of eukaryotic algae. Like other important transcription factors, the gene encoding RpaB cannot be deleted, making the study of deletion mutants impossible. Based on a bioinformatic approach, we increased the number of known genes controlled by RpaB by a factor of 5. Depending on the distance to the TSS, RpaB mediates transcriptional activation or repression. The high number and functional diversity among its target genes and co-regulation with other transcriptional regulators characterize RpaB as a regulatory hub.

## INTRODUCTION

RpaB (“regulator of phycobilisome association B”) is an OmpR-type transcription factor of crucial importance for the transcriptional control of multiple photosynthesis-associated genes under varying light conditions. RpaB was discovered in *Synechocystis* sp. PCC 6803 (from here: *Synechocystis* 6803) based on its ability to affect the energy distribution from phycobilisomes (PBS) to photosystem I (PSI) relative to photosystem II (PSII) (1). Orthologs of *rpaB* belong to the cyanobacterial core genome and were discovered in the chloroplast genomes of all non-green algae except *Odontella* (2) and in the charophyte alga *Chlorokybus atmophyticus* (see (3) for a recent review), suggesting important and evolutionarily widely conserved functions. The *rpaB* gene cannot be deleted by conventional methods, indicating its essentiality (1, 4–7). Despite its wide distribution, most insight into the genes controlled by RpaB have been obtained from analyses in two model cyanobacteria, *Synechocystis* 6803 and *Synechococcus elongatus* PCC 7942 (from here: *S. elongatus*).

The RpaB binding motif consists of a pair of imperfect 8-nt long direct repeats (G/T)TTACA(T/A)(T/A) separated by two random nucleotides. This motif was found upstream of many genes responding to high light (HL) in both *Synechocystis* 6803 and *S. elongatus* and was named the HLR1 (“high light regulatory 1”) sequence (4, 8). The binding of RpaB to the HLR1 sequence was first demonstrated for the *hliB* gene promoter in *Synechocystis* 6803 (9). Binding to HLR1 promoter elements of the *hliA* and *hliB* genes encoding HL-inducible proteins leads to repression under LL, both in *Synechocystis* 6803 and *S. elongatus* (6, 9, 10). Chromatin immunoprecipitation analysis showed that the binding activity of RpaB to the *hliA* and *rpoD3* promoters in *S. elongatus* was promptly lost upon a shift to HL (11), leading to de-repression of these HL-inducible genes. The same mode of transcriptional regulation was reported for the sRNA gene *psrR1* in *Synechocystis* 6803 (12). For several genes encoding PSI proteins, binding of RpaB to HLR1 was identified to be crucial for the transcription activation under low light (LL) (13, 14). Chromatin affinity purification (ChAP) analysis revealed that loss of binding activity of RpaB upon the shift to HL leads to a large decline in the transcript levels of PSI genes, while expression of PsrR1 is induced (12). PsrR1 is a negative post-transcriptional regulator of genes encoding phycobiliproteins and subunits of PSI (15), leading to the dual repression of PSI genes under HL - at the transcriptional level by RpaB and at the post-transcriptional level by PsrR1 (12). The effect of RpaB binding on target promoters, which may be activation or repression under LL, is likely determined by the location of the HLR1 motif. However, some of the mentioned genes underlie complex transcriptional controls, including multiple transcriptional start sites (TSSs) belonging to separate promoters. This is the case for the *psaAB* dicistron, for which three separate TSSs and three distinct HLR1 elements have been detected in *Synechocystis* 6803. Of these, two activate and one represses transcription under LL, and it is the joint regulation at these three sites that leads to the observed regulation (14). In *S. elongatus*, in addition to photosynthesis-related genes, RpaB binds to the promoters of the sigma factor genes *rpoD3* and *rpoD6,* and also the promoter of the core circadian clock genes *kaiBC* (16, 17). To date, 83 occurrences of the HLR1 motif have been reported including examples from several different cyanobacteria, the *Cyanophora* chloroplast and cyanophage genomes (3).

Therefore, it is established that the regulon controlled by RpaB consists of genes encoding photosynthesis-related proteins as well as at least one sRNA and that this regulation is of crucial importance in light acclimation responses and circadian clock-related processes (18). In contrast, its cognate histidine kinase Hik_33_ (synonyms NblS or DspA) functions as multistress sensor responding not only to HL (19) but also to low temperature (20), hyperosmolarity (21), high salinity (22), oxidative stress (23) and nutrient stress (7). Therefore, it is likely that only a small portion of the RpaB functions has been discovered thus far. In particular, a global definition of the RpaB regulon is missing.

Here, we combined existing data from the genome-wide high-precision mapping of TSSs (24–26) with the precise identification of regulated promoters, *in silico* motif prediction, functional enrichment analysis and validation experiments to infer the RpaB regulon in a comprehensive way, choosing *Synechocystis* 6803 as a model.

## RESULTS

### HLR1 elements are predicted at two distinct positions relative to the TSS

Based on the sequence alignment of 90 previously reported HLR1 motifs associated with 83 promoters in *Synechocystis* 6803, *S. elongatus*, other cyanobacteria and eukaryotic algae (3), a position specific weight matrix (PSWM) was generated (**Figure S1**), which served as input for the global motif search, as outlined in **Figure 1**. We analyzed the positional distribution of 3,615 theoretically possible HLR1 sites (**Table S1**) in the promoters of 1,992 transcriptional units (TUs) in *Synechocystis* 6803. HLR1 elements clustered at two distinct sites, centered ~51 nt upstream of the TSS or overlapping the TSS (**Figure 2A**). The precise positions for these two enriched HLR1 occurrences were from −66 nt to −45 nt (peaking at position −51) and from −38 nt to +23 nt (centered at position −5) relative to the respective TSS. By comparing the respective promoters against comparative primary transcriptome data (24), these two distinct peaks matched different expression profiles. Genes with an HLR1 at or close to the TSS were generally more upregulated under HL, whereas genes with an HLR1 at −51 were generally more downregulated under HL (**Figure S2**). Therefore, the peak around −51 was identified as belonging to HL-repressed/LL-activated promoters, whereas the motifs centered at −5 belong to HL-activated/LL-repressed promoters (**Figure 2B**). This is consistent with previous reports showing that binding of RpaB to HLR1 more distally located to the TSS is crucial for activating transcription under LL (13, 14), whereas derepression under HL, e.g., of PsrR1, was connected to an HLR1 motif overlapping the TSS (12). However, here this observation is extended to a large set of promoters. Hence, depending on the distance between the HLR1 sequence and the TSS, RpaB can be either stimulating or repressing (see also (3, 27) for review). For clarity, we will speak of genes activated or repressed by RpaB under LL when referring to these regulatory phenomena in the remainder of the manuscript.

**Figure 1.**
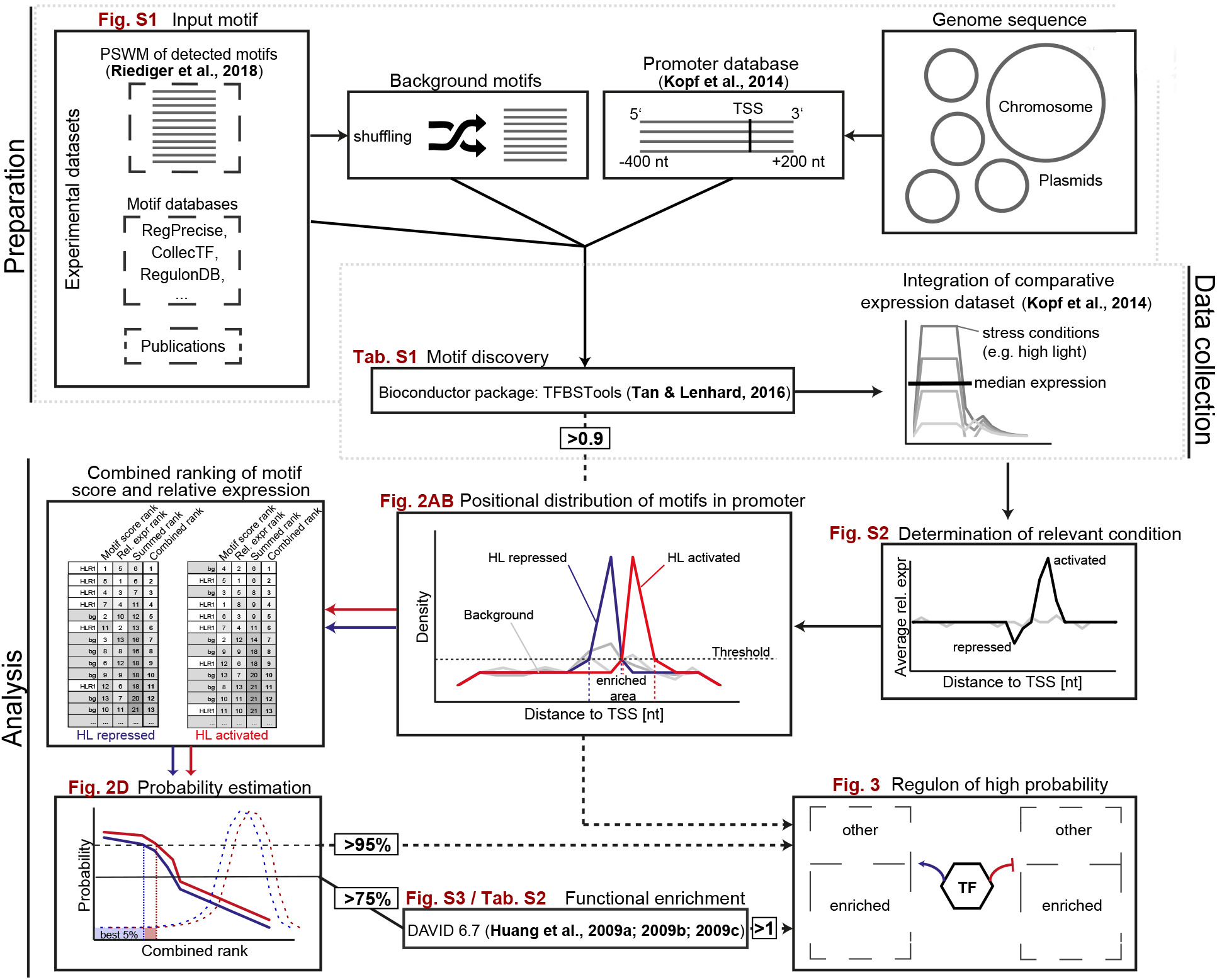
Bioinformatic workflow. **Preparation:** A *Synechocystis* 6803 promoter database was created by extracting sequences from −400 nt to +200 nt relative to the transcriptional start sites (TSSs) of all transcriptional units (TUs) in 5‘ to 3‘ orientation as identified by Kopf et al. (2014). The HLR1 input motif was defined as a position specific weight matrix (PSWM, **Figure S1**) via the alignment of all published HLR1 sequences (3). The background motifs were generated by permuting the columns of the PSWM. **Data collection:** Motif detection was performed using the Bioconductor package “TFBSTools” (70). The comparative dRNA-seq gene expression dataset by Kopf et al. (2014) was integrated with the results, and the relative expression under each of the ten tested conditions was normalized against the median expression of the gene of interest. **Analysis:** The average relative expression level of each promoter with an HLR1 was plotted against the relative distance to the TSS to determine conditions correlating with a particular location. High light (HL) was identified as a relevant condition for further analysis (**Figure S2**). Subsequently, the positional distribution of the HLR1 within the promoters was examined (**Figure 2A**). Motifs whose promoters were activated or repressed under the relevant condition were separated prior to this analysis and the density of detected motifs in activated or repressed promoters was plotted against their relative distance to the TSS (**Figure 2B**). Areas exceeding a density threshold of ≥ 99% were considered significantly enriched, and only motifs occurring in one of these areas were used for further analysis. Motifs from each enriched area were ranked according to their motif score and relative expression and probabilities were calculated from the overall distribution of the motifs’ combined ranks (**Figure 2D**). A probability threshold of ≥ 75% was set to perform a functional enrichment analysis of all genes meeting this criterion (**Figure S3, Table S2**) by using DAVID 6.7 (29–31). All genes in a group with an enrichment score ≥ 1, a probability ≥ 95% or a relative motif score ≥ 0.90 were accepted for the final regulon (**Figure 3**). The regulon was visualized using Cytoscape 3.5.1 (76).

**Figure 2.**
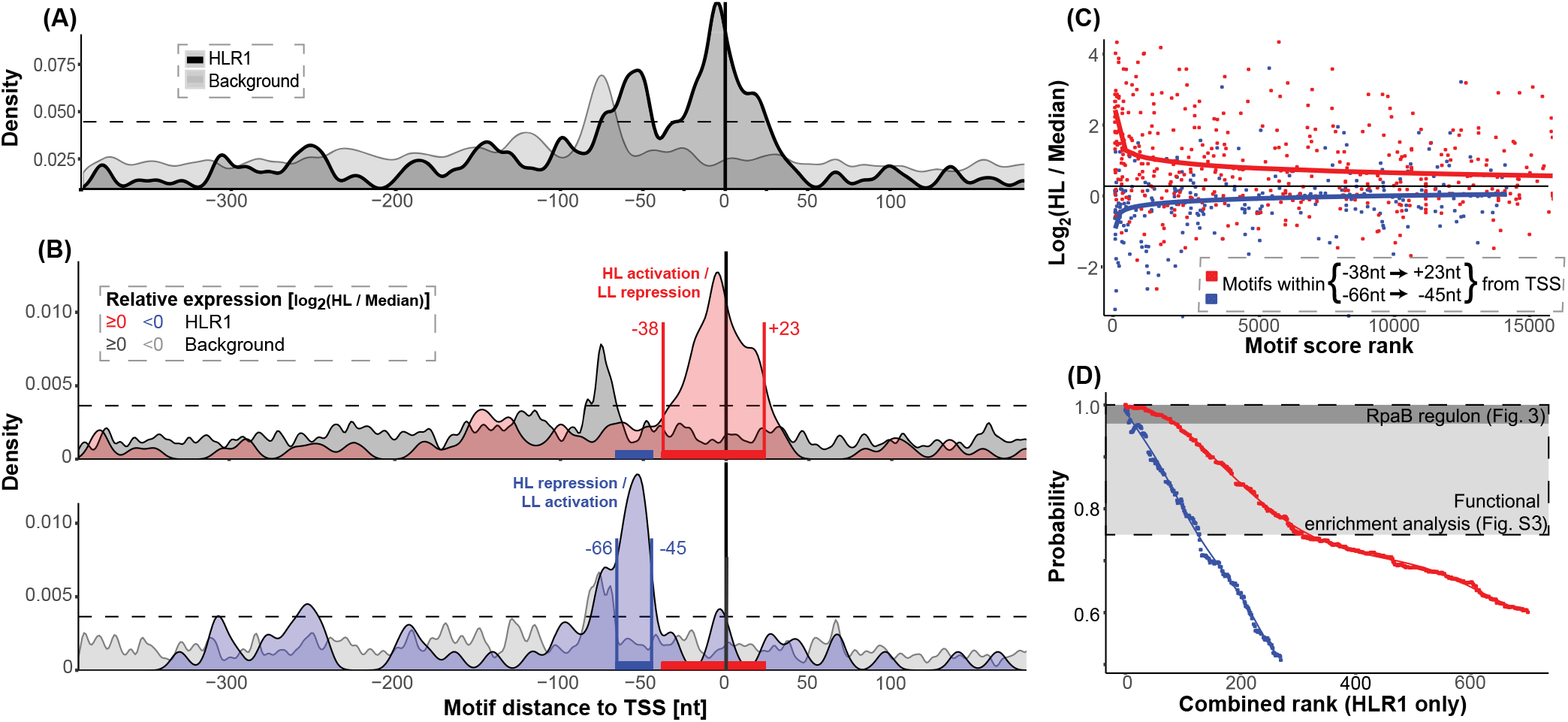
Bioinformatic analysis of HLR1 occurrences in *Synechocystis* 6803. **(*A*)** Positional distribution of HLR1 motifs within the set of promoter sequences. A background model was created by permuting (n = 100) the PSWM columns of HLR1 (**Figure S1**) and detecting occurrences of these random motifs in all promoters of *Synechocystis* 6803 (binwidth = 5 nt; relative motif scores ≥ 0.90). Throughout the analyses, the same parameters were applied for the background model as well as for HLR1. **(*B*)** Two distinct locations of the HLR1 motif correlate with activation or repression under HL (**Figure S2**). Promoter sequences were separated prior to this analysis according to their response to a shift to HL [log_2_(HL / median) ≥ 0 or < 0], their density was plotted against their relative distance to the TSS (binwidth = 5 nt; relative motif scores ≥ 0.90) and enriched areas were defined (density threshold ≥ 99%). The quality of the HLR1 elements from each enriched area was rated by ranking their relative motif scores (output from TFBSTools) against the set of generated background motifs (total rank) and the correlation between the relative expression under HL. **(*C*)** Ranking the relative expression against the HLR1 motif score. A combined rank was calculated for each motif (HLR1 + background), and probabilities were obtained from the distribution of the motifs’ combined ranks. **(*D*)** Comparison of the combined HLR1 ranks against their respective probabilities.

Interestingly, similar peak patterns as those observed under HL were detected for cold (15°C) and –Fe conditions (**Figure S2**), which is consistent with the fact that the RpaB/Hik33 two-component system acts as a redox sensor for several external stimuli and controls the electron transfer routes, especially in cold-signal transduction (28).

### The combination of computational motif detection, dRNA-seq expression data and functional enrichment analysis reveals a large and complex RpaB regulon

Three criteria were used to filter the HLR1 occurrences in the two distinct peak areas: (i) a probability threshold ≥ 95%, (ii) a relative motif score ≥ 0.90, and (iii) a functional enrichment score > 1. At least one of these criteria had to be met for inclusion, yielding a list of 98 activated genes (under control of 43 promoters) and 196 repressed genes (under control of 124 promoters) (**Table S3**).

The ranked motif scores correlated well with the relative expression of the associated promoters under HL (**Figure 2C**). Accordingly, the combined rank of motif score and relative expression allowed the computation of a probability for each HLR1/target to be regulated by RpaB (**Figure 2D**). The probability estimation was generated from the distribution of the background.

Functional enrichment analyses were conducted for the filtered TUs and those down to a probability ≥75% (cf. **Figure 2D**) using DAVID 6.7 (29–31) (**Figure S3**, **Table S2**). Targets that were predicted to be repressed by RpaB were enriched in genes with a wide range of functionalities, such as nitrate assimilation, circadian clock, GTP-binding, PSII reaction center subunits and components of associated photosynthetic electron transport chains (**Figure S3A**). In contrast, genes predicted to be activated by RpaB were enriched for encoding subunits of PSI or the PBS (**Figure S3B**) but also included also *isiA* and *lexA* and, with the CRISPR2 system, one of the three native CRISPR-Cas systems present in this organism (32).

Using conservative parameter settings, the final set of putative RpaB targets encompassed 188 promoters (**Table S3, Figure S4**). Of these, 166 control the expression of protein-coding genes and 22 control non-coding TUs. The predicted regulon included 36 of 39 genes or operons previously shown to be under RpaB control in *Synechocystis* 6803 (**Figure S4**), which validated the performance of our approach. Hence, these results raised the number of putative RpaB targets in *Synechocystis* 6803 by approximately 5-fold for all promoters. The three genes or operons that were not identified either lacked a TSS in our dataset (*slr0897*), the reported motif differed too much from the HLR1 model employed here (*petE/sll0199*), or the distance to the TSS was higher than that allowed in this analysis (*psbH/ssl2598*). Putative targets were not restricted to the chromosome but were also found on plasmids, suggesting the deep integration of those genes in the regulatory network.

To gain more information about different functionalities, we split the full set of predicted target genes into subclusters (**Figure 3**) based on functional enrichment analysis. Genes that are important for photoprotection (encoding HLIPs, carotenoid biosynthesis, glutathione/glutaredoxin), photorespiration or synthesis of C1 compounds and cofactors (*gcvP)* photosynthetic electron transport (*petF1-3*, *petE*, *pgr5)*, hydrogenase formation (*hypD/slr1498*), cofactor biosynthesis (heme & porphyrin, quinones, riboflavin), nitrogen assimilation and the circadian clock were restricted to the repressing branch of RpaB, consistent with the higher demand for these activities under HL. Additionally, many genes encoding stress-related regulatory proteins such as proteases, RNases, chaperones and GTPases were predicted repressed targets, including three of the four genes encoding subunits of FtsH protease complexes.

**Figure 3.**
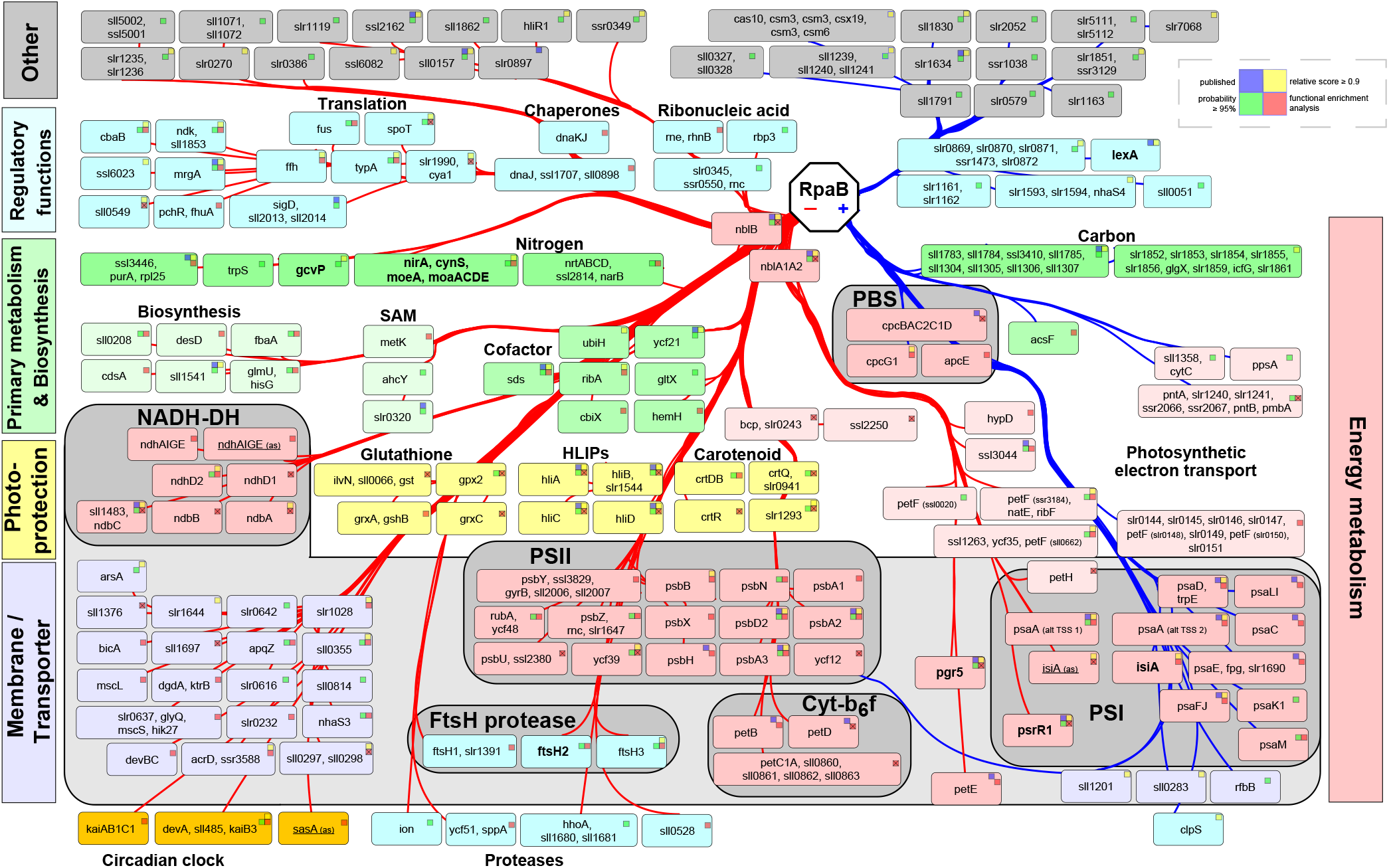
RpaB regulon of high probability. All RpaB targets detected by at least one criterion are displayed. A set of up to 4 squares indicates if the regulation was previously published (blue square), if the probability was ≥ 95% (green square), if the relative motif score was ≥ 0.90 (yellow square) and a functional enrichment score of ≥ 1 (red square). A + sign within the squares indicates manual curation due to incomplete database annotation. Putative ncRNAs / asRNAs that have not been further described in the literature are not displayed. The targets were arranged according to their predicted regulation by RpaB and colored according to their membership in functional groups or complexes (for further information about the individual target genes see **Table S3**). Underlined targets indicate asRNAs.

### A specific view on photosynthesis-related genes reveals the antagonistic regulation of most PSII and cytb_6_f versus PSI genes by RpaB binding to different HLR1 positions

With few exceptions, most PSII genes possess an HLR1 within or close to the area where RpaB acts as a repressor, suggesting coordinated co-downregulation via RpaB (**Figure 4**). Similar observations were made for several genes of the cytochrome-b_6_f complex and soluble electron acceptors (such as *petE*, *pgr5*, *petF*). On the contrary, most PSI genes harbor an HLR1 in the area where RpaB acts as an activator (**Figure 4**). A few examples, for which regulation cannot be intuitively inferred, possess complex promoter organization, such as in the case of *psaA* (14), or multiple instances of an HLR1 motif, as for *psbA1*, *psbA2* and *psaM* (**Figure 4**). The *isiA*-IsrR pair is very intriguing: IsrR is the asRNA controlling the accumulation of *isiA* mRNA (33). Both promoters have an identical HLR1 element. However, this element is in a repressing position for IsrR and an activating position in the *isiA* promoter.

**Figure 4.**
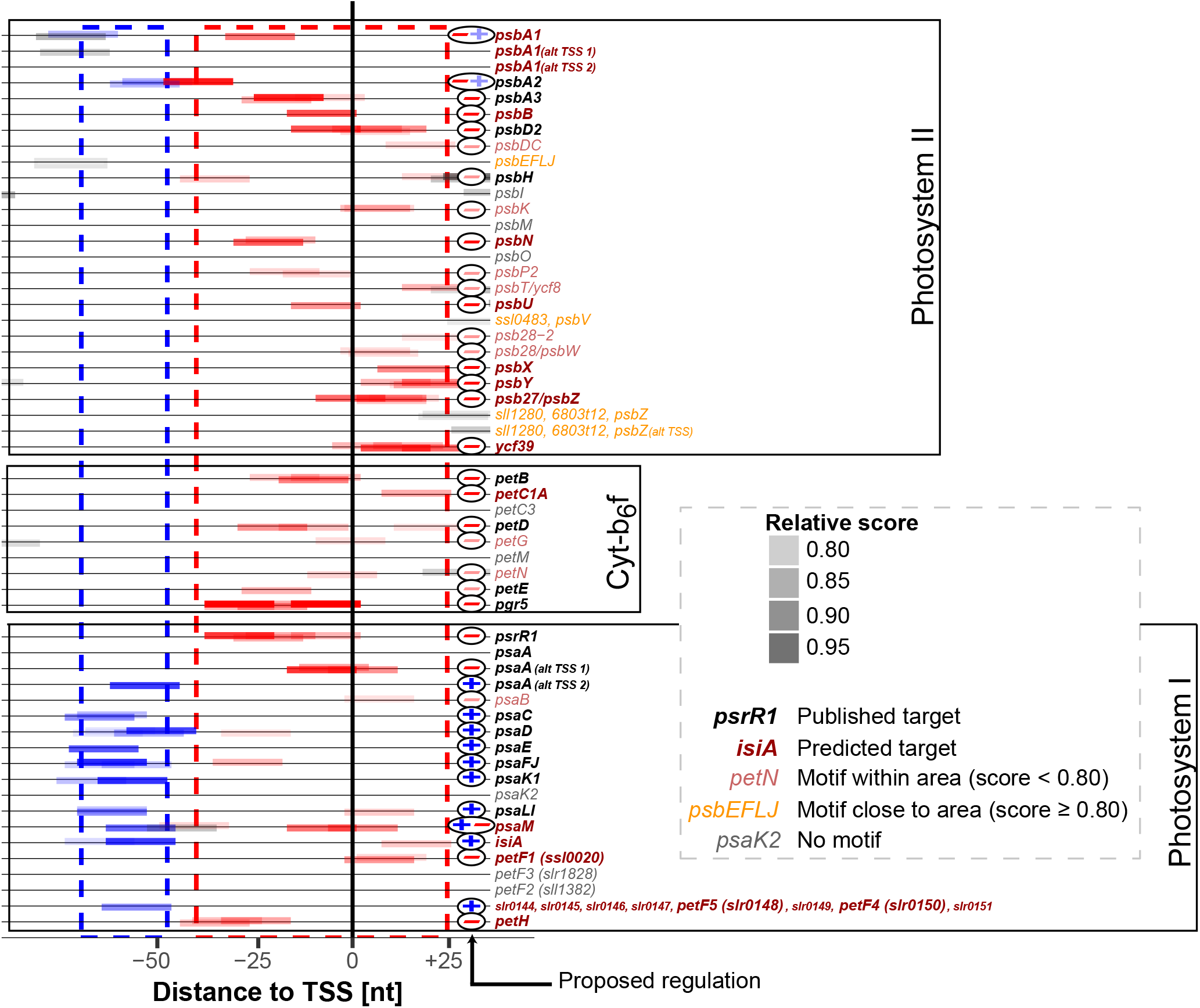
HLR1 motifs within promoters of the photosynthetic apparatus. Detected HLR1 motifs (score ≥ 0.75) were plotted against their distance to the corresponding TSS and colored according to the deduced regulation (blue = activation by RpaB; red = repression by RpaB under LL; gray = regulation was not deduced because motif was outside of the relevant two regions). The saturation of the motif reflects the consensus with the input motif. The prominent + or – symbols indicate the resulting regulation.

### Novel members of the RpaB regulon stimulated by HL have functions in cyclic electron flow, photorespiration and nitrogen assimilation

The *pgr5* gene is strongly induced when cells are exposed to HL or to CO_2_ limitation (24). This gene encodes a homolog of PGR5 (proton gradient regulation 5) in *Synechocystis* 6803 (34), which is involved in antimycin A-sensitive electron flow from PSI to the plastoquinone pool in plants and algae (35, 36). The *pgr5* promoter possesses twin HLR1 boxes, HLR1a and HLR1b, covering the −35 to −10 elements and the first transcribed nucleotide (**Figure 5A**). ChAP assays utilizing 12xHis-tagged RpaB from a LL-grown culture yielded 32% recovery of this promoter fragment, while this number dropped to ~5% at 5 min after transfer to HL and recovered to only 11% at 15 min after the shift, which was even lower than the negative control (**Figure S5**). In promoter fusion experiments, the native P_*pgr5*_ promoter mediated a rapid induction of luciferase fluorescence upon the shift from LL to HL. This promoter derepression was consistent with the rapid induction of *pgr5* mRNA accumulation in the wild-type (WT) 5 min after the shift from LL to HL, shown here by Northern blot hybridization and in microarray expression data taken from the CyanoEXpress database (37). Mutation of HLR1b abolished reporter gene expression, while mutation of HLR1a alone or of both motifs together led to an more pronounced peak upon transfer from LL to HL (**Figure 5A**), pointing to additional regulatory mechanisms. It should be noticed that *pgr5* was the most differentially regulated gene in a recently found regulatory mechanism involving the protein Slr1658 (38). Therefore, EMSAs were performed, which showed direct binding of RpaB to the native sequence. Binding was still detectable with individual HLR1a or HLR1b mutants but completely abolished when HLR1a and HLR1b were jointly replaced (**Figure 5A**).

**Figure 5.**
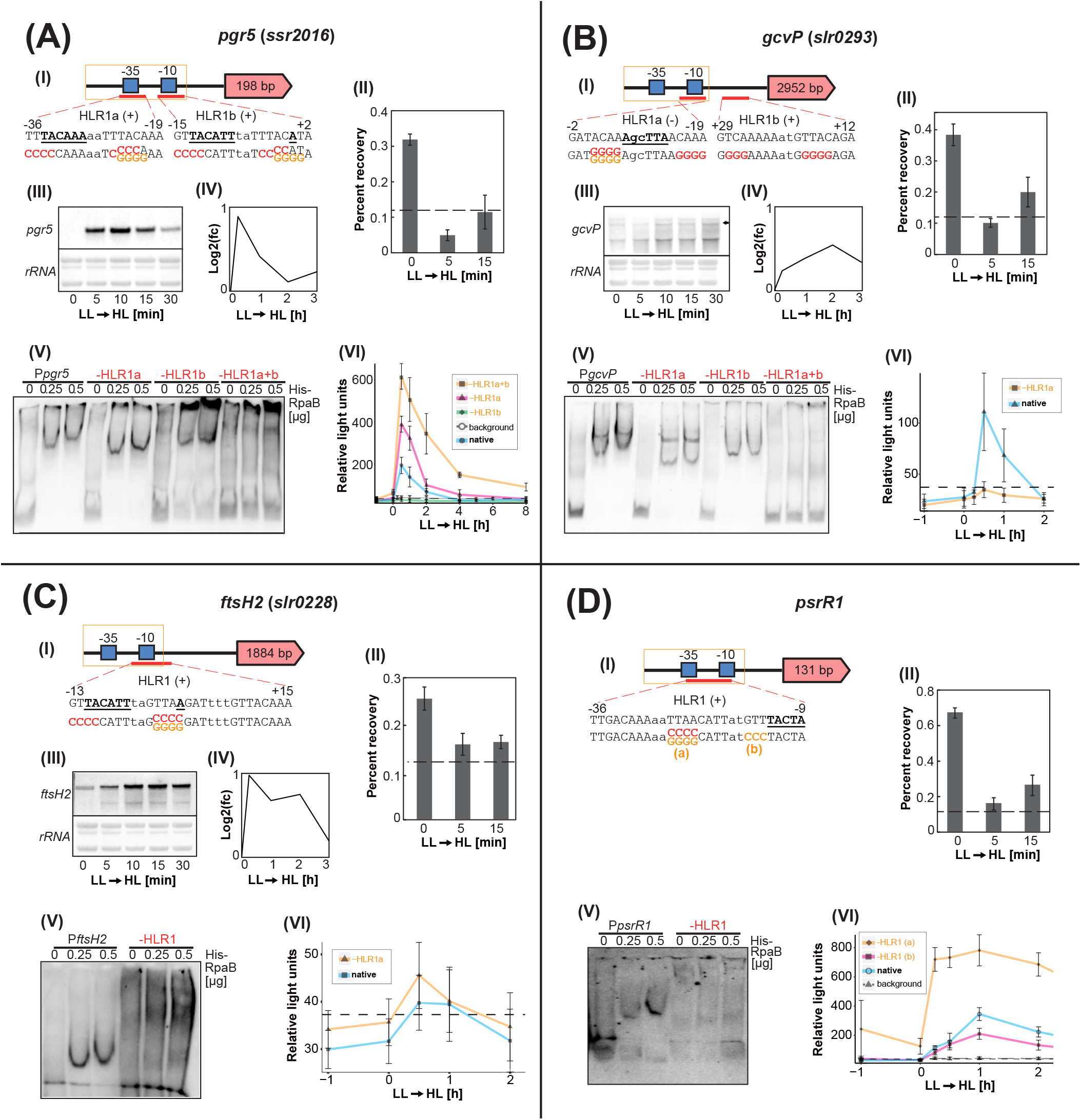
Experimental validation of selected potential target genes predicted to be repressed by RpaB. **(*A***) *pgr5 (ssr2016).* **(*B***) *gcvP (slr0293).* **(*C***) *ftsH2 (slr0228). psrR1* serving as a positive control (12) for the conducted experiments**. (I) Schematic representation of individual promoters.** Red bars give the HLR1 location, (+) or (-) signs indicate the orientation on the forward or reverse strand. The sequences are given in capital letters underneath, other promoter elements and the TSS are underlined and in boldface. Introduced point mutations are marked in red for the EMSA (III) and in orange for the luciferase assay (VI). **(II) ChAP analysis.** Percent recovery after the shift to HL. The dashed line resembles the maximum percent recovery which was obtained from the negative control (**Figure S5**). Error bars represent means ±SD from three independent experiments. **(III) Northern blot.** Transcript levels after the shift to HL. **(IV) Expression profile.** Relative expression after shift to HL, taken from the CyanoEXpress database (37). **(V) Electrophoretic mobility shift assays (EMSA).** Binding of RpaB to the native and mutated promoters. The sites of the introduced mutations are depicted in red in the HLR1 shown in (I). **(VI) Luciferase reporter assays.** Absolute luminescence [relative light units] under HL exposure (alternatively, nitrogen depletion + HL exposure in the case of *nirA*). Error bars represent the standard deviation from triplicates of two biological replicates and from at least three independent experiments. The native promoters that were used are boxed in orange in part (I). The background luminescence (average background = dashed line) originated from a strain containing only the decanal-producing plasmid, used as a negative control.

The *slr0293/gcvP* gene encodes the P-protein, one of four subunits of the glycine decarboxylase complex (39) involved in the plant-like photorespiratory C2 cycle metabolizing poisonous 2-phosphoglycolate, which is a by-product of the bifunctional Rubisco activity. Therefore, the enhanced expression of *gcvP* with increased irradiance is sensible and was observed by Northern blot hybridization here as well as in microarray expression data from the CyanoEXpress database (**Figure 5B**). The *gcvP* promoter possesses twin HLR1 boxes. HLR1a covers the −10 element and the region up to −2 of the TSS, while HLR1b is located within the 5’UTR (**Figure 5B**). ChAP assays using RpaB from a LL-grown culture yielded 38% recovery of this promoter fragment. This number dropped to ~10% at 5 min after transfer to HL and recovered to 22% at 15 min after the shift. In promoter fusion experiments, the native P_*gcvP*_ promoter mediated an induction in luciferase fluorescence upon a shift from LL to HL within 30 min. Joint mutations of HLR1a and HLR1b abolished reporter gene expression, consistent with EMSA data demonstrating the lack of RpaB binding to the fragment when both sites were mutated (**Figure 5B**).

The FtsH2 protease, which is important for PSII repair (40) is also controlled by RpaB. P_*ftsH2*_ possesses a single HLR1 overlapping the TSS and the −10 element (**Figure 5C**), suggesting a repressing function under LL. Indeed, Northern hybridization demonstrated the induction of *ftsH2* mRNA accumulation in the WT upon the shift from LL to HL, but the initial level was higher than for *pgr5*. ChAP assays yielded 27% recovery of this promoter fragment, while this number dropped to ~15% at 5 min after transfer to HL, below the negative control value obtained with the *glnB* promoter (**Figure S5**), suggesting actual RpaB binding. Binding was directly confirmed in EMSA assays with the native promoter sequence, while it was abolished when the motif was mutated (**Figure 5C**). Reporter gene assays confirmed the inducibility by HL. The HLR1 mutation led to reduced, but still detectable, HL-inducible expression of the reporter gene. We conclude that P_*ftsH2*_ is under RpaB control and that the existing 2^nd^ TSS secured the lower, albeit still detectable, regulation when HLR1 was mutated.

We chose the promoter of PsrR1 as a positive control (12, 15). P_PsrR1_ possesses an HLR1 spanning the region from the −35 to the −10 element (**Figure 5D**). Indeed, ChAP assays yielded 65% recovery of this promoter when protein extracts from LL-grown cultures were applied, while this number dropped to less than 20% 5 min after transfer to HL and recovered to 25% at 15 min after the shift. Moreover, EMSAs showed direct binding of RpaB to the native sequence, while binding was not observed when the HLR1 motif was mutated. In promoter-reporter gene fusion assays, we observed rapid de-repression upon the shift to HL and even stronger derepression when the HLR1 motif was replaced. In contrast, mutation of the −10 element led to low expression and almost no detectable light induction (**Figure 5D**). These facts corresponded well to the known regulation of this sRNA (15).

### HLR1 motifs in complex promoter architectures suggest the integration of redox signaling with different environmental signals transmitted by other transcription factors

Several putative RpaB binding sites were detected in promoters that are known to be regulated by other transcription factors such as Fur, LexA or NtcA. Hence, the promoters of the *isiA*, *nirA* and *lexA* genes were examined to gain insight into the cross-regulation between RpaB and other transcription factors. The gene *isiA* encoding the chlorophyll-binding protein CP43′ or IsiA (<u>i</u>ron <u>s</u>tress <u>i</u>nduced protein A) is repressed under iron-replete growth conditions by the Ferric uptake regulator Fur (41) and it is de-repressed under iron starvation (42, 43). The *isiA* promoter was predicted with high confidence as an RpaB target. The HLR1 motif finishes only 5 nt upstream of one of the two Fur-binding elements (**Figure 6A**). RpaB binding was confirmed by the ChAP assay; also recovery of the native promoter fragment dropped 5 min after the shift to HL. Binding of both transcription factors, RpaB as well as Fur, to the expected promoter motifs was confirmed by EMSA. Although the HLR1 motif was identified in the activating position, we could not detect *isiA* transcripts under LL conditions (**Figure 6A (III), +Fe**). However, after incubation under iron-deplete conditions for 48 h, high transcript accumulation was detected, with a slight decrease after the shift to HL (**Figure 6A (III), 48 h-Fe**). The reason for this finding is it’s parallel repression by Fur, as validated by the reporter gene assays. When the cells were starved for iron, gene expression from the native promoter increased linearly for 36 h, whereas mutation of the Fur box caused very high expression before iron was removed, consistent with the role of Fur as a repressor. However, when HLR1 was mutated, no expression could be detected at all, even if the Fur box was mutated in parallel. This result suggested that RpaB was required for the high-level expression of *isiA* under iron-limiting conditions. Consistent with this interpretation, the shift from LL to HL led to a remarkable decrease in gene expression with both the native and Fur box-mutated promoter variants, followed by an increase when the cells were shifted back (**Figure 6A (VI)**).

**Figure 6.**
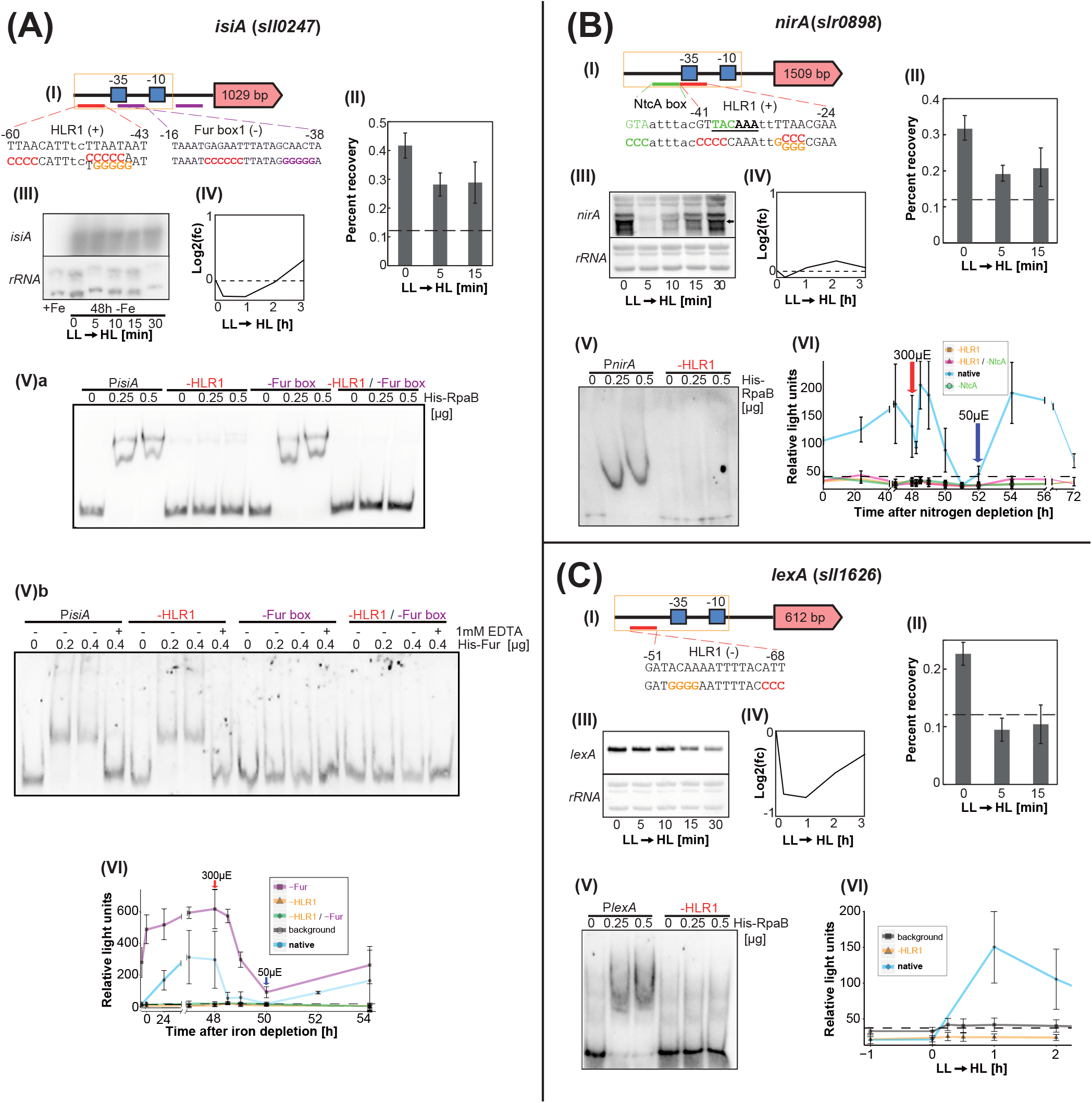
Experimental validation of selected potential RpaB target genes showing cross-regulation with other transcriptional regulators. **(A**) *isiA (sll0247).* The Fur boxes were taken from publication (50). **(*B***) *nirA (slr0898).* The NtcA box was taken from publication (44). **(C**) *lexA (sll1626)*. **(I-VI)** Refer to explanations in **Figure 5**.

The *nirA* gene encoding nitrite reductase belongs to the core regulon controlled by global nitrogen regulator NtcA (44). The NtcA box within P_nirA_ is overlapping the HLR1 by 5 nt (**Figure 6B (I)**). Under nitrogen-replete growth conditions, *nirA* transcript levels declined immediately after exposure to HL, then increased to the level higher than that under LL (**Figure 6B (III and IV)**). Such tendency was also observed in promoter activity under nitrogen-depleted conditions (**Figure 6B (VI)**). Point mutations into either HLR1 or NtcA boxes resulted in decreased luminescence relative to the natural promoter (**Figure 6B (VI)**). EMSAs showed a simple band pattern and specific binding of RpaB to the HLR1 sequence in P_*nirA*_, whereas binding was abolished when HLR1 was mutated.

LexA is a highly conserved transcription factor throughout the bacterial domain and frequently functions as a repressor of SOS response-related genes involved in DNA repair (45). In *Synechocysti*s 6803, however, LexA has been reported to act as a global regulator binding to the promoters of genes contributing to various cellular functions, such as hydrogenase activity (46), bicarbonate transport (47), twitching motility (48), glucosylglycerol accumulation (48) and fatty acid biosynthesis (49). The promoter of *lexA* harbors a single HLR1 in the activating position (**Figure 6C (I)**). Our experimental data show that RpaB bound to P_*lexA*_ with extracts from LL-grown cells, while this binding was lost 5 min after transfer to HL, and mutation of HLR1 within P_*lexA*_ prevented RpaB binding (**Figure 6C (II and V)**). Moreover, fusion of the native promoter to the reporter did indeed yield fluorescence, while HLR1 mutation prevented it. Hence, activation of P_*lexA*_ by RpaB under LL was validated. Consistently, decreasing *lexA* transcript levels were observed after transfer to HL (**Figure 6C (III and IV)**). However, *lexA* promoter activity increased after 1 h of HL exposure (**Figure 6C (VI)**). This increase in promoter activity was not observed when a mutation was introduced into HLR1 (**Figure 6C (VI)**), indicating that this increase was due to the re-binding of RpaB to HLR1. Although LexA has been suggested to be negatively auto-regulated (49), our results did not show any clue for self-repression of the *lexA* gene.

## DISCUSSION

### Extension of the RpaB regulon

Our results substantially increase the number of known RpaB-regulated promoters in *Synechocystis* 6803, from 37 to at least 167 for protein-coding genes or operons and from 1 to 22 non-coding RNAs. To infer the regulon controlled by RpaB, we analyzed the positional distributions of RpaB binding sites, the expression profiles of the regulated genes and performed functional enrichment analysis. For seven selected examples, we performed reporter gene assays, ChAP and EMSA assays and measured mRNA accumulation by Northern hybridizations. The results from the luciferase reporter gene assays revealed inducibility as early as 30 min after exposure to HL for promoters that were predicted to be repressed by RpaB under LL (*ftsH2*, *pgr5*, *gcvP* and *psrR1* for control), hence confirming the validity of our approach.

### RpaB as a key regulator for redox regulation and acclimation to HL

Several new putative target genes associated with photosynthetic functions or electron transfer were identified. These findings add to the known role of RpaB as a key regulator of many essential subunits of the photosynthetic apparatus and strongly support that RpaB is involved in redox regulation and acclimation to HL. Our data demonstrate the involvement of RpaB in the control of *pgr5/ssr2016*, which is involved in cyclic electron flow, and in the expression of *gcvP/slr0293*, encoding the glycine decarboxylase P-protein, one of the four subunits of the glycine decarboxylase complex. The finding of dual HLR1 boxes in the *pgr5/ssr2016* promoter is intriguing. The twin HLR1 elements likely ensure tight regulation under LL, similar to the role of the twin Fur boxes in the *isiA* promoter (50). The encoded Pgr5 protein is involved in cyclic electron flow from PSI to the PQ pool in cyanobacteria, plants and algae alike (34). The major function of cyclic electron flow is to balance ATP/NADPH ratios and preventing PSI photoinhibition (51). Therefore, it makes perfect sense that *pgr5* is controlled by RpaB. However, PSI photoinhibition is also prevented by Mehler-like reactions mediated by the Flv1 and Flv3 proteins (52), although the genes encoding these proteins (*sll1521* and *sll0550*) were not found here, suggesting a more specific role for RpaB in state transition and cyclic electron flow. Further predicted target genes encode components of the NADH dehydrogenase complex such as *ndhD2/slr1291* and *ndhAIGE*(*sll0519-sll0522*), consistent with the suggested link between these complexes and cyclic electron transport (53–55). Other functionally connected targets involve subunits of the cytochrome-b_6_f complex, plastocyanin (*petE*), five ferredoxins (*ssl0020*, *sll0662*, *ssr3184*, *slr0148*, *slr0150*) (fed1-5 = all plant-like ferredoxins), as well as *petH*, encoding the ferredoxin-NADP(+) oxidoreductase, bacterioferritin (Bcp/Slr0242) and its associated ferredoxin (Ssl2250), which were reported to be involved in resistance to oxidative stress by utilizing thioredoxin as a reductant (56). Thioredoxin was shown to directly influence the RpaB redox state (57), and therefore RpaB might indirectly control its own activity via the downregulation of *bcp* and *ssl2250*. Altogether, RpaB controls six out of nine ferredoxins present in *Synechocystis* 6803 (58). These findings illustrate the importance of regulation of the electron distribution within and downstream of photosynthesis via RpaB.

Photorespiratory 2-phosphoglycolate (2-PG) metabolism is essential in cyanobacteria and especially important during HL acclimation (59, 60). The production of 2-PG promotes the accumulation of glycine in HL, which is toxic in high concentrations (61). Therefore, the glycine cleavage complex is important for HL acclimation processes as well. Redox regulation of *gcvP,* encoding the P-protein subunit of this complex, was reported (62), which, according to our data, is performed by RpaB. As proposed for P*pgr5*, the possession of two HLR1 elements might ensure a tight regulation. The cellular antioxidant defense system consists of various mechanisms to maintain redox homeostasis, which is crucial to cope with stress conditions, such as HL. RpaB seems to regulate most genes of the glutathione/glutaredoxin system, relevant to prevent photooxidative damage. GrxA and GrxC were reported to be the only HL-inducible glutaredoxins, serving as electron acceptors of glutathione (63).

### Cross-regulatory effects between RpaB and other regulatory mechanisms

Our results for the genes *isiA* and *nirA* point to cross-regulation with the transcription factors NtcA and Fur, respectively.

Mediated by Fur, many cyanobacteria respond to iron starvation by expressing IsiA (42, 43). IsiA initially forms antenna rings around trimeric PSI (64) and likely serves as an extra light-harvesting complex, whereas it functions later in the dissipation of excess light energy (65, 66) and exerts a protective function (67). Repression of *isiA* by Fur under iron-replete growth conditions was validated by the high luciferase expression measured when the Fur box was mutated and the fact that the native promoter became induced 24 h after the removal of iron ions (**Figure 6A (VI)**). In contrast, mutagenesis of the HLR1 site completely abolished *isiA* expression, even under iron-deplete conditions, demonstrating the function of RpaB as an activator of *isiA* gene expression. Our identification of RpaB as an activator of *isiA* gene expression adds up to the tight repression by Fur and the asRNA IsrR (33), which was predicted to be repressed by RpaB at LL (**Figure 3**, **Table S3**). Therefore, the here suggested involvement of RpaB in the transcriptional regulation of *isiA* and, inversely in the control of its asRNA IsrR, adds a regulatory dimension that has been previously missing.

RpaB control over the glutathione/glutaredoxin system illustrates its interface with nitrogen metabolism, which is important for enhancing the electron sink capacity. Cross-regulation with NtcA in the control of *nirA/slr0898,* encoding ferredoxin-nitrite reductase, also fits this picture. Binding of RpaB to the *nirA* promoter was clearly shown; however, the HLR1 is at an intermediate position at −41 ~ −24, which is still compatible with activation by RpaB. Therefore, the observed decline after HL shift in the amount of the *nirA* transcript in the presence of combined nitrogen (**Figure 6B (III) and (IV)**) and in the promoter activity in the absence of combined nitrogen (**Figure 6B (VI)**) can be explained by the release of RpaB. An overcompensation observed after initial decline was likely mediated by NtcA. Upon the inverse shift from HL to LL, the observed increase in activity was solely due to activation by RpaB. Hence, the *nirA* promoter is co-stimulated by NtcA and RpaB. However, an asRNA overlapping the first codons and 5’UTR of *nirA* has been detected (24, 44) and may contribute to the control of mRNA stability and the efficiency of translation in the native situation.

Further predicted targets encode proteins involved in HL acclimation, such as PSII subunits or *slr0228* and *slr1604* encoding the FtsH2/3 protease complex, which is important for PSII repair (40). It is intriguing that the FtsH2 protease is also required for the induction of inorganic carbon acquisition complexes in *Synechocystis* 6803, acting upstream of the transcription factor NdhR (68), and that the FtsH1/3 heterocomplex controls the level of Fur (69). All three genes (*ftsH1, ftsH2, ftsH3*) are repressed by RpaB (**Figure 3**).

Together with previous data, the results of this work show that RpaB is of crucial and central relevance for the light and redox-dependent remodeling of the photosynthetic apparatus, various electron transfer chains, state transitions and, overall, for the light acclimation of photosynthetic cyanobacteria. The RpaB regulon encompasses many additional genes that are more loosely or not at all associated with the photosynthetic machinery, exemplified by three of four genes encoding FtsH heterocomplexes, or the glutathione/glutaredoxin system. RpaB is involved in the control of carbon and nitrogen primary metabolism, the circadian clock and exerts cross-regulation with other transcription factors such as NtcA or FurA. The size and importance of the controlled regulon characterize RpaB as one of the most crucial transcriptional regulators in cyanobacteria.

## MATERIALS AND METHODS

### Bioinformatics Methods

A workflow illustrating the major bioinformatics steps, tools, important data and results is given in **Figure 1.**

A *Synechocystis* 6803 promoter database was set up and the HLR1 motif defined as presented in **Figure 1.** Motifs were detected with the Bioconductor package TFBSTools (70) was used (relative motif score ≥ 0.8), and integrated withdifferential RNA-Seq data obtained previously (24). The average relative expression of all promoters with an HLR1 at the same relative position was plotted against the relative distance to the TSS and further examined as outlined in **Figure 1.**

### Experimental Methods

#### Strains and growth conditions

*Synechocystis* 6803 was grown in BG11 (71) buffered with 20 mM N-[Tris(hydroxymethyl)methyl]-2-aminoethanesulfonic acid (TES), pH 7.6, at 30°C at 50 μmol quanta * m^-2^ * s^-1^ of white light under slight agitation ± selective antibiotics (kanamycin, Km: 50 μg*mL^-1^, chloramphenicol, Cm: 20 mg*mL^- 1^). To induce HL stress, cultures were shifted from 30 to 300 μmol quanta * m^-2^ * s^-1^. To induce iron deficiency, 300 μM desferrioxamin B (DFB) was added to liquid cultures. To induce nitrogen deficiency, cells were centrifuged, washed and cultivated in NO_3_^-^- free BG11.

#### Construction, expression and purification of *E. coli* strains expressing His-tagged transcription factors

The coding region of *furA* (*sll0567*) was amplified by PCR using the primers *Nde*I-FurA-fw and *Xho*I-FurA-rv (oligonucleotides are listed in **Table S4**) and cloned into the pT7Blue T-vector (Novagen). The *furA* PCR fragments were excised from the pT7Blue vector with *Nde*I and *Xho*I and subcloned into the same restriction sites in vector pET28a r (Novagen) to express proteins with an N-terminal 6xHis-tag. Following transformation into *E. coli* ArcticExpress (DE3)RIL competent cells (Agilent), the strains harboring the Fur expression construct were precultured in 5 mL LB medium containing 50 μg*mL^-1^ kanamycin at 37°C overnight. The preculture was seeded into 500 mL LB and FurA expression was induced with 100 μM IPTG from midlog cultures grown overnight at 15°C. Purification of His-Fur proteins was performed on a nickel column using the ÄKTA start system (GE Healthcare). The purity of His-FurA was checked by SDS-PAGE. The preparation was desalted by PD MidiTrap G-25 (GE Healthcare) and concentrations were determined via Bradford assay. The construction, expression and purification of *E. coli* strains expressing His-tagged RpaB was performed as described (12).

#### ChAP

Preparation of whole-cell extracts for ChAP analysis, affinity purification of His-RpaB and DNA were performed as described (12) with some modifications as follows. A 0.5 mg aliquot of protein of the whole-cell extract and 10 μL of Dynabeads His-Tag Isolation and Pulldown (Veritas) were used for affinity purification of His-RpaB, followed by further purification steps and quantitative real-time PCR analysis as described (12).

#### Northern blot

Isolation of RNA by the hot-phenol method and RNA gel blot analyses using the DIG RNA labeling and detection kit (Roche) were performed as described (12). To facilitate the direct use of PCR products as templates for *in vitro* transcription, the T7 polymerase promoter (TAATACGACTCACTATAGGGCGA) was added at the 5**’** termini of the reverse primers.

#### Electrophoretic mobility shift assay (EMSA)

The *nirA* and *ftsH2* promoter fragments, with or without mutations, were PCR-amplified from *Synechocystis* genomic DNA using Go Taq Hot Start Green Master Mix. Native and mutated promoter regions of *pgr5*, *gcvP*, *psrR1* and *lexA* were generated by annealing overlapping primers and filling up without template DNA. Native and mutated versions of the *isiA* promoter were amplified from the pILA vector containing this promoter (72). The DNA fragments were 3’ end-labeled with digoxigenin (DIG)-ddUTP by terminal transferase according to the manufacturer’s instructions (DIG gel shift kit; Roche) with some modifications to remove EDTA using NucleoSpin Gel and PCR Clean-up (Macherey-Nagel). Assays were performed as described (12, 73).

#### Luciferase assay

Promoters containing predicted motifs of interest were amplified, encompassing ~400 nt upstream of the TSS, and introducing *Age*I and *Fse*I restriction sites at the 5’ and 3’ ends, respectively. PCR products were double-digested, ligated with the linearized empty pILA vector and transformed into chemically competent *E. coli* DH5α cells. To introduce site-directed mutagenesis, non-overlapping primers containing the desired mutation were designed by using the NEBaseChanger web tool (http://nebasechanger.neb.com/), followed by inverse PCR. The generated amplicons were *Dpn*I digested, phosphorylated, self-ligated and transformed into chemically competent *E. coli* DH5α cells. Following DNA preparation and cleavage by *Age*I and *Fse*I, the promoters of interest were inserted upstream of the *luxAB* 5’UTR in plasmid pILA (72). The *Synechocystis* 6803 luciferase reporter strains harbored the decanal-producing plasmid pTCmYfr2luxCDE containing a *luxCDE* cassette with the Cm marker under control of the *yfr2a* promoter (74, 75). A strain harboring only the decanal-producing plasmid served as a negative control. Liquid precultures (two biological replicates per strain) were diluted (OD_750_ = 0.1) prior to analysis and cultivated to exponential phase (OD_750_ = 0.6 – 0.8). Prior to measurement, cells were diluted to an OD_750_ = 0.6, aliquots of 100 μL were transferred in three technical replicates per strain to a 96-well microtiter plate (PerkinElmer) and luminescence was measured as reported (74).

## ACKNOWLEDGEMENTS

We thank Martin Hagemann, University of Rostock, for critical reading of the manuscript.

## Supplementary Figure legends

**Figure S1.**
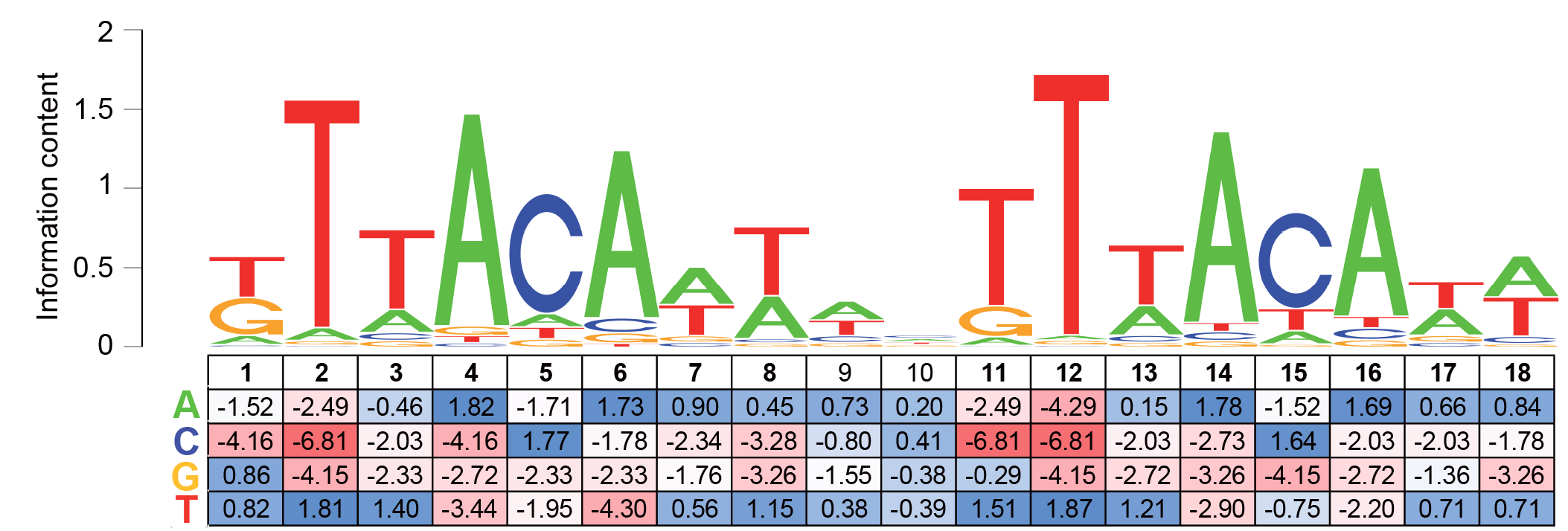
Position specific weight matrix (PSWM) of HLR1 used as input for the global motif search. HLR1 from 90 previously described motifs found in 83 promoters in *Synechocystis* 6803, *S. elongatus*, other cyanobacteria and eukaryotic algae (3).

**Figure S2.**
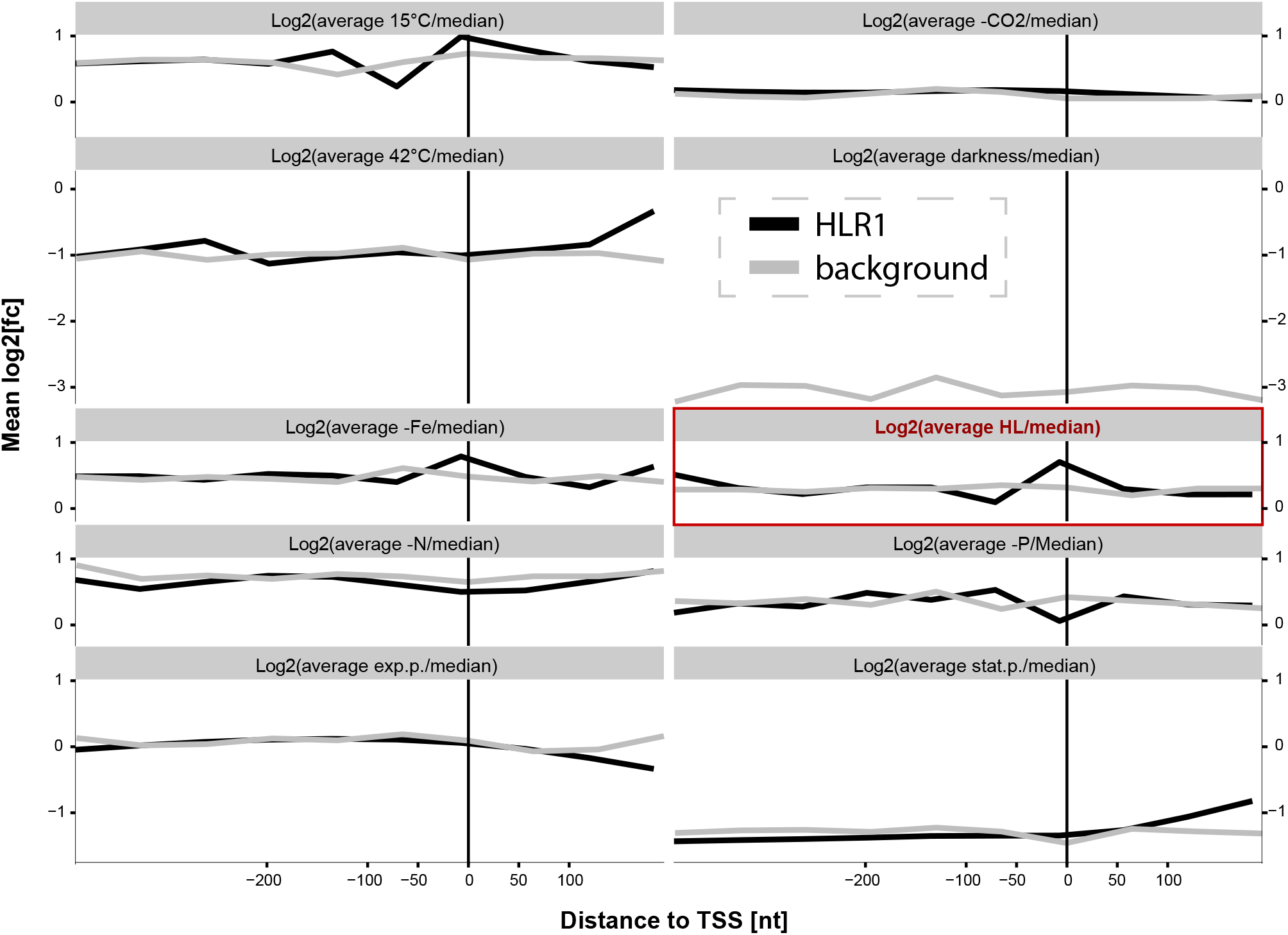
Average relative expression (condition / median expression) of all genes having an HLR1 at the same relative position within their promoters plotted against the distance of the motif to the TSS. The data were taken from the ten different conditions investigated in the differential RNA-Seq analysis by Kopf et al. (24). The same analysis was performed with the background model as a negative control.

**Figure S3.**
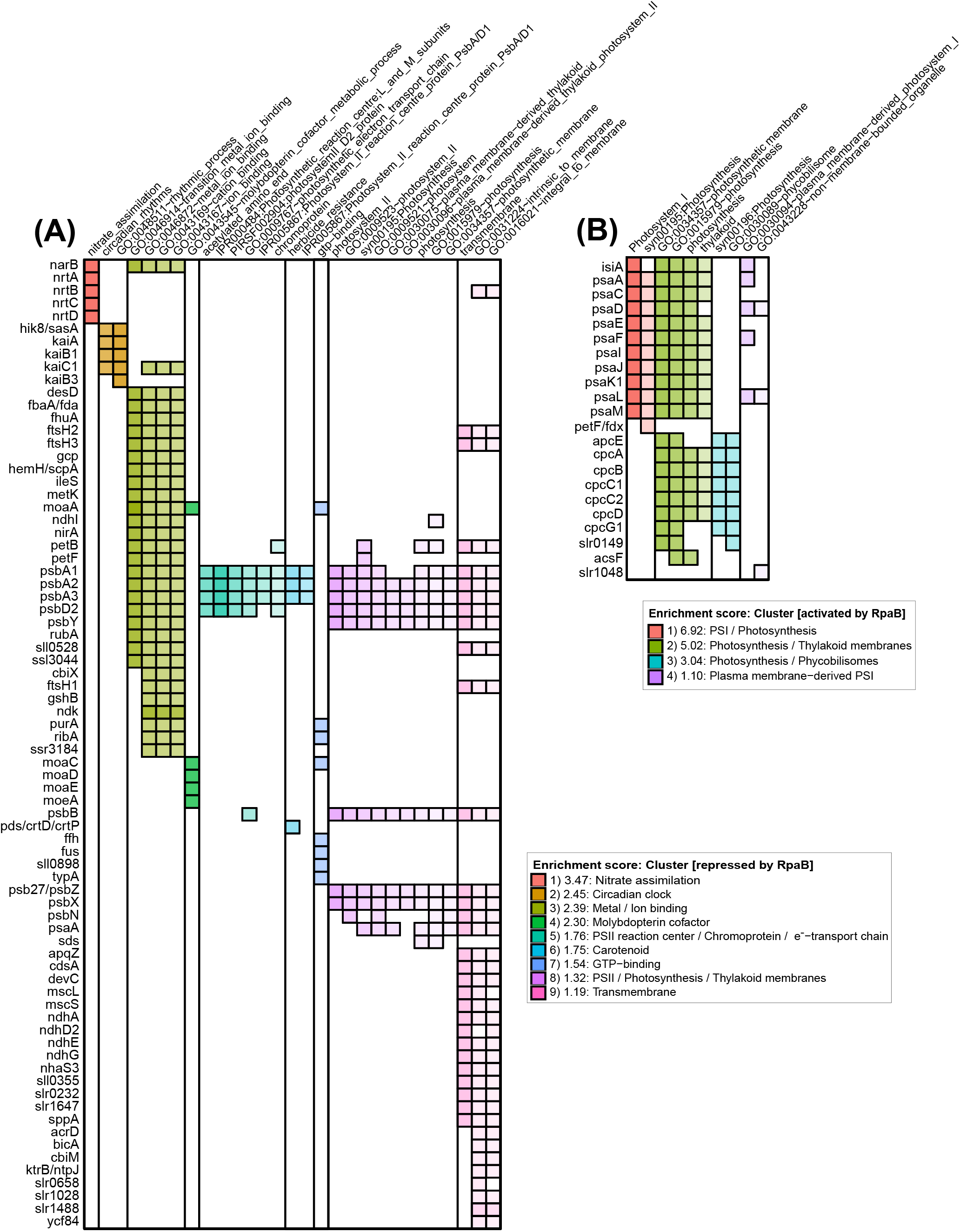
Functional enrichment analysis. Gene clusters which were predicted to be **(*A*)** repressed or **(*B*)** activated by RpaB are shown in a heat map. A probability threshold ≥75% was set to perform the functional enrichment analysis by using DAVID 6.7 (29–31). All clusters with a highly stringent enrichment score ≥ 1 are shown (see also **Table S2** for further information).

**Figure S4.**
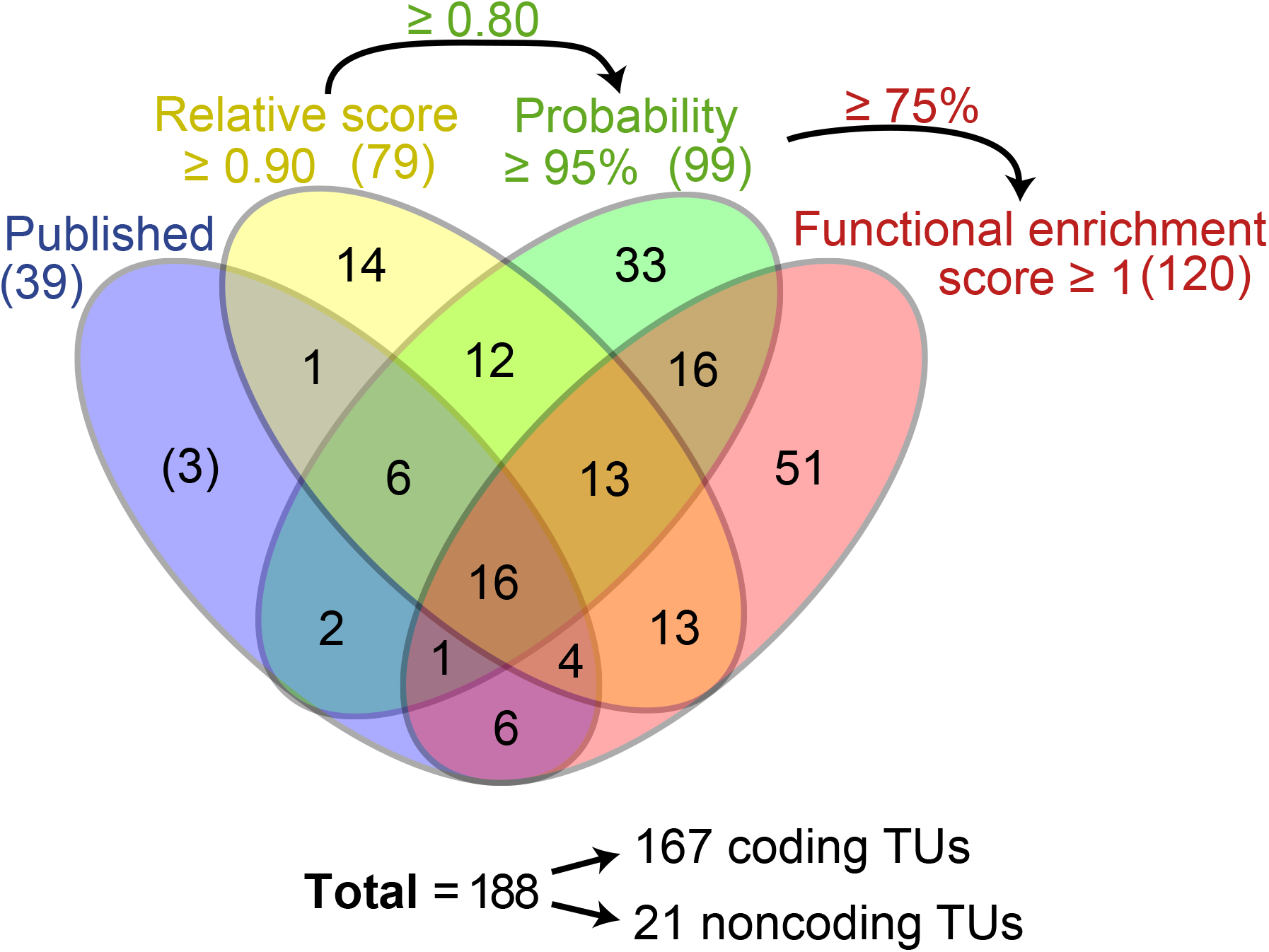
Venn diagram showing the number of genes or operons detected here by the combination of the three given parameters compared to the number of previously published targets in *Synechocystis* 6803. The selected parameters for the further analyses (motif score ≥ 0.80, probability ≥ 75%) and for the acceptance for the final regulon (motif score ≥ 0.90, probability ≥ 95%, functional enrichment score ≥ 1) as well as the total number of coding and non-coding TUs of the final regulon with these parameter settings are given.

**Figure S5.**
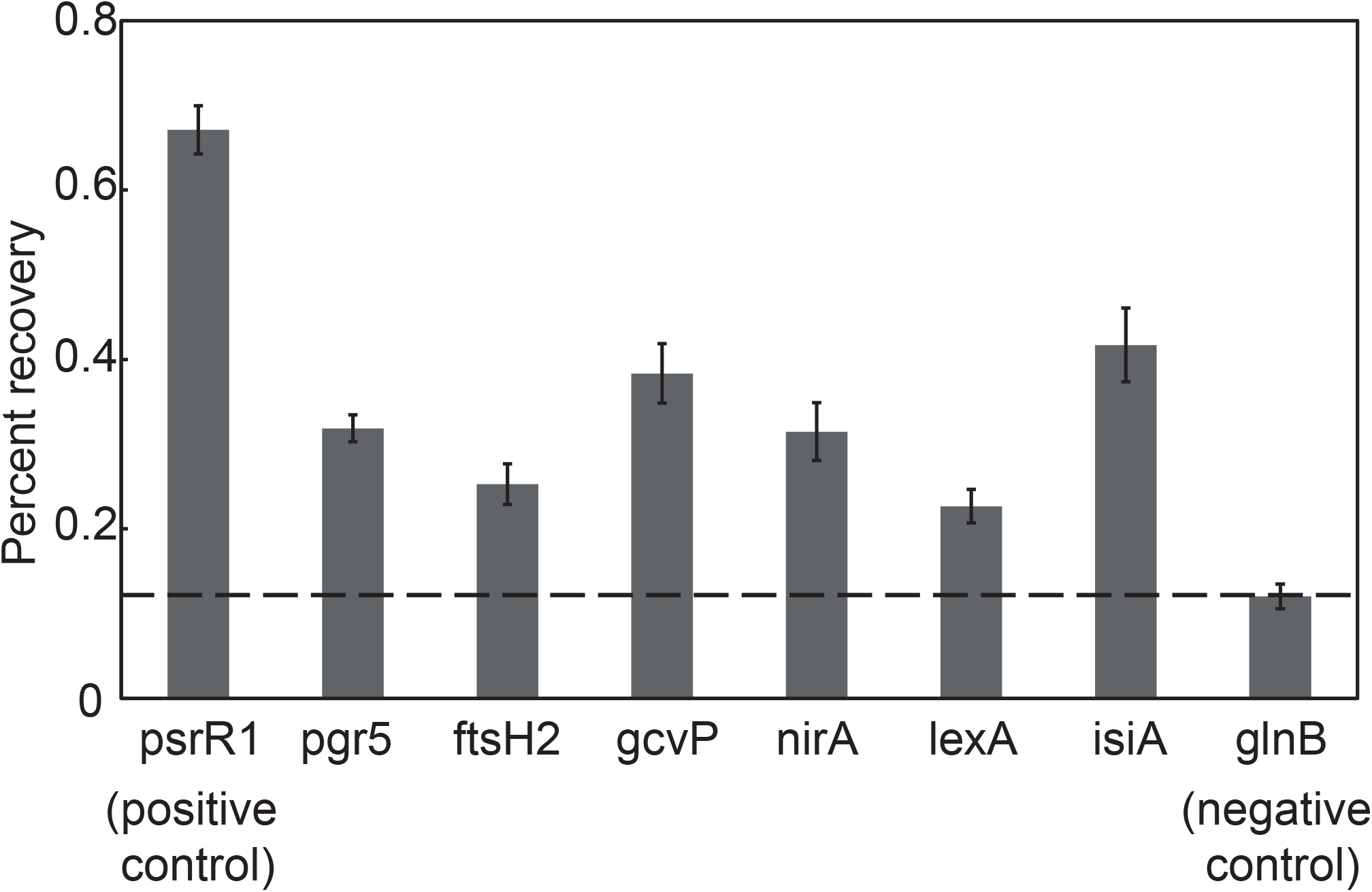
Binding activity of RpaB to the putative target promoters under LL conditions examined by ChAP (negative control: *glnB*). The level of DNA co-purified by nickel chromatography was determined by qPCR analysis and is given as percentage recovery relative to the total input DNA. Data are means ±SD from three independent experiments.

**Figure S6.**
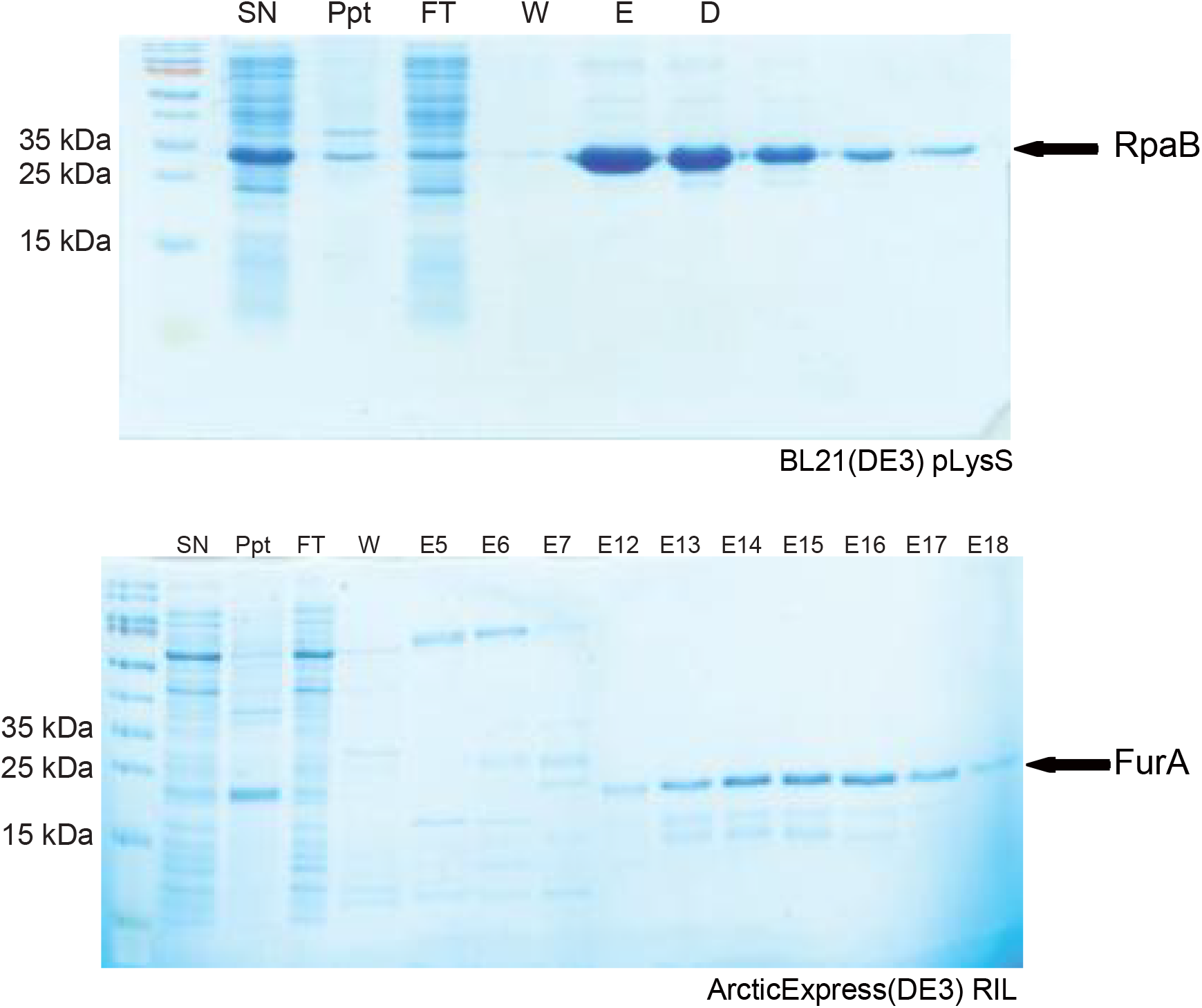
Coomassie gel showing the purification of His-RpaB and His-FurA by nickel affinity chromatography. The soluble fraction from an *E. coli* over-expression strain was loaded onto a HiTrap chelating HP column (GE Healthcare). Aliquots of supernatant (SN), insoluble precipitation (Ppt), flow-through (FT), wash (W) and eluate fractions were separated by SDS-PAGE and stained with Coomassie Brilliant Blue.

**Table S1-S4:** Attached as one Excel file

**Supplementary Table legends**

**Table S1.** Raw output from R-package TFBSTools.

**Table S2.** Functional enrichment analysis.

**Table S3.** All predicted RpaB targets and HLR1 sites. Information if the respective promoter was previously identified as an RpaB target in Synechocystis 6803, in another species or was chosen for detailed experimental validation is given in columns AG to AI.

**Table S4.** Desoxyoligonucleotide sequences.

